# Receptor-structured modelling of EGFR-driven tumor initiation: from spatially resolved cell-based simulations to reduced population dynamics

**DOI:** 10.64898/2026.06.05.730529

**Authors:** Romasa Qasim, Anass Bouchnita

## Abstract

Alterations in epidermal growth factor receptor (EGFR) dynamics can influence tumor initiation by changing receptor abundance, ligand-dependent activation, and downstream proliferative signaling. Mathematically linking these receptor-scale processes to population-level tumor growth remains challenging because they couple molecular, cellular, and tissue-scale dynamics. Here, we develop multiscale models that explicitly captures receptor-ligand dynamics. We analyze the dynamics of a refined version of a 3D stochastic multicellular model with explicit EGFR-EGF interactions to derive a receptor-structured continuum model in which cells are organized by active receptor clusters. This model is further reduced into a population dynamics model that tracks the mean number of active receptors. It captures the main qualitative behaviours of the higher-dimensional models while enabling analytical and numerical characterization of model-derived thresholds for sustained growth. After calibration and comparison with available *in vivo* tumor-growth data under EGFR overexpression, we use the model hierarchy to quantify how initiation thresholds depend on EGF availability, EGFR abundance, receptor-ligand unbinding, and genetic potential. The models predict that EGFR overexpression, stronger receptor-ligand binding, and more aggressive cell phenotypes each lower the EGF molecular counts required for sustained tumor growth. Overall, the proposed framework provides a flexible mathematical approach for connecting receptor-ligand kinetics with population-level tumor-initiation dynamics.

## 1 Introduction

Epidermal Growth Factor Receptor (EGFR) is a Receptor Tyrosine Kinase (RTK) that forms clusters [1] and is often overexpressed in various cancers such as breast, glioma, ovarian, non-small-cell lung, prostate, pancreatic, and head and neck cancers [2], and correlates with progression and poor prognosis. Mutant forms of EGFR, such as the constitutively active EGFRvIII, contribute to cancer heterogeneity. EGFR plays a critical role in maintaining cancer stem cells (CSCs), supporting their stemness, metabolism, immunomodulation, dormancy, and resistance to therapy [3]. Aberrations in the EGFR pathway hyperactivate their G-protein domains and downstream pro-oncogenic pathways, such as RAS-RAF-MEK-ERK MAPK and AKT-PI3K-mTOR [4]. These pathways drive cancer cell proliferation by promoting chronic activation of transcription factors through the cell cycle, as shown in Figure 1 [5]. Moreover, EGFR is commonly overexpressed and causes resistance to different treatments such as radiation therapy [6] and chemotherapy [7]. Further, EGFR overexpression is used as a predictive biomarker for the efficacy of tyrosine kinase inhibitor (TKI) treatments [8]. These studies highlight the importance of developing a deeper understanding of how changes in EGFR expression and EGFR-EGF binding dynamics impact tumor initiation, progression, and treatment.

**Fig 1.**
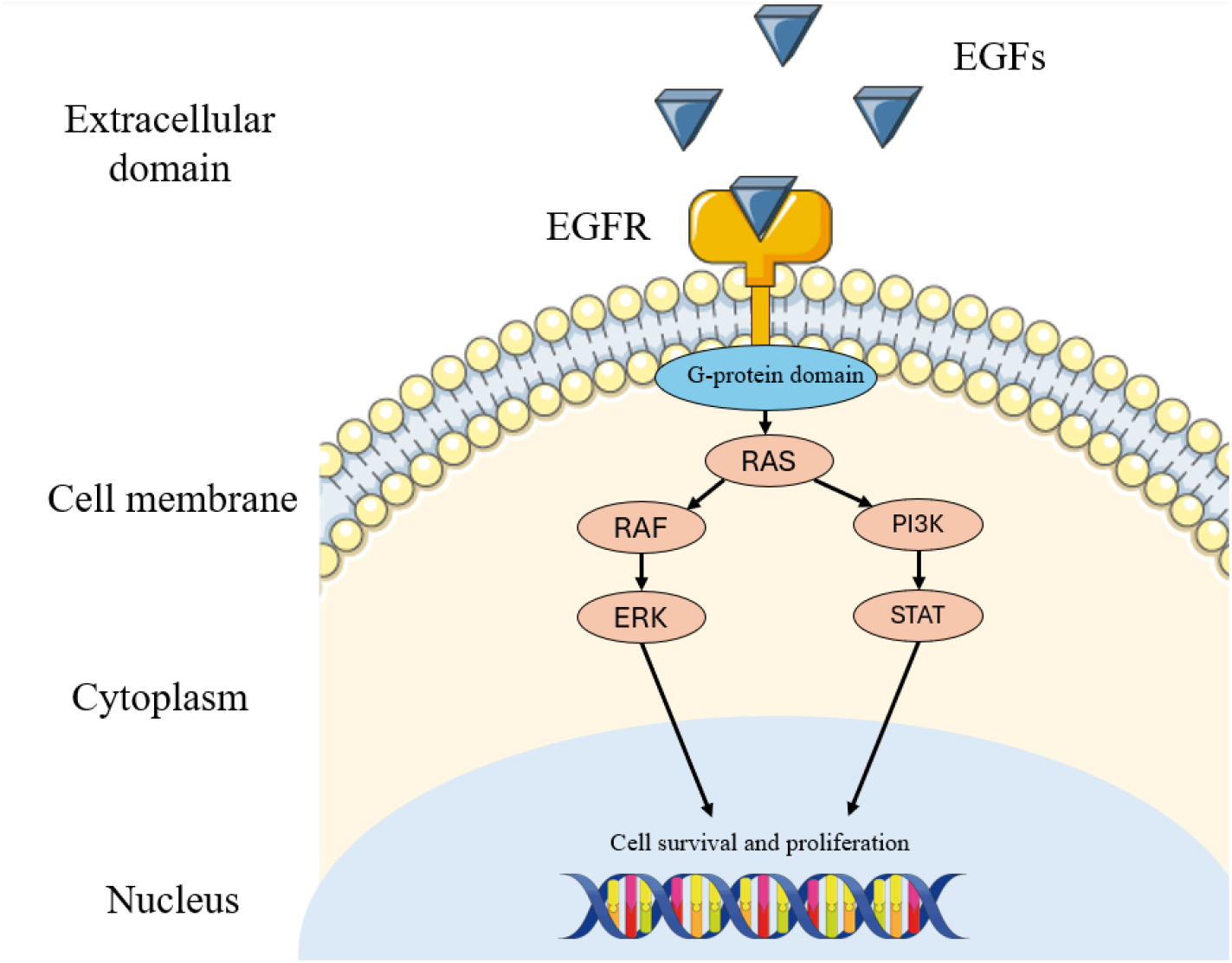
A schematic representation showing the interaction between the epidermal growth factor (EGF) and its receptor (EGFR) as well as the downstream activation of the ERK and PI3K pathways, which promote cell survival and proliferation. After their binding to EGF molecules, active EGFR molecules phosphorylate their G-protein binding site, located on the cytoplasmic side of the cell membrane. These sites activate the RAS molecules, which turn on the ERK and PI3K molecules. These molecules activate transcription factors that promote cell survival and proliferation, leading to tumor initiation.

Epidermal growth factor (EGF) is an agonist ligand that binds to an enzyme-linked receptor, specifically the epidermal growth factor receptor (EGFR), a receptor tyrosine kinase. Similar to other receptor-ligand connections, binding affinity is a key metric for describing biomolecular interactions and often serves as a surrogate for activation efficacy. It describes the ability of the ligand to bind for a long time to receptors. Many methods have been developed to calculate receptor-ligand binding affinities, including molecular dynamics (MD) [9, 10], Brownian dynamics (BD) [11], and Markov state modelling [12]. Mechanistic mathematical models, commonly formulated as a set of ordinary differential equations, have provided valuable insights into ligand-receptor binding dynamics, building on foundational biochemical principles developed over the past century [13, 14]. Several models used more complex stoichiometric systems to study questions such as the transition between various receptor states [15, 16], impact on intracellular regulation [17, 18, 19], and integration with pharmacokinetics-pharmacodynamics (PK-PD) [20].

Cancer arises from excessive cell division caused by acquired mutations. These mutations disrupt intracellular signaling, enhancing cell survival and proliferation. Continuous, discrete, and multiscale mathematical models have been developed to gain insight into complex tumor cell dynamics [21, 22]. Population dynamics models have been extended to study key biological processes regulating tumor growth, such as growth types [23], [24], angiogenesis [25], immune response interactions [26, 27], extracellular cytokine effects [28], and treatment [29]. Further, they can be expanded to include spatial effects such as the ones caused by cell motility or cytokine diffusion [30]. Several models of this class were developed in 1D, 2D, and 3D to describe the process of malignant cell invasion [31, 32], the spheroid structures of tumors [33, 34], and the interplay between cancer and tumor cells [35, 36]. Other extensions have been developed to capture tumor cell evolution during tumor progression by stratifying cells according to their phenotype [37, 38, 39]. In many population-dynamics models, receptor activation is represented through an effective growth rate. This abstraction is useful for capturing population-level behavior, but it does not separate specific receptor-level processes such as ligand-receptor association, dissociation, binding affinity, or agonist competition.

Discrete models of tumor growth represent individual cells and their interactions using methods such as cellular automata, on-lattice, or off-lattice approaches [40, 41, 42, 43, 44, 45, 46]. These models provide valuable insights into tumor emergence and growth by focusing on cell-cell interactions, which improves their ability to represent complex tumor dynamics [47, 45]. Hybrid models combine discrete cell representations with continuous equations for intracellular and extracellular dynamics [48, 49]. Hence, they integrate the strengths of both approaches [50, 51, 52, 53]. Further, they allow the incorporation of data collected from different approaches such as imaging, flow cytometry, and transcriptome sequencing. Hybrid models typically describe intracellular and extracellular protein concentrations with differential equations while representing cells as discrete objects. As a result, they capture cell spatial heterogeneity and genotype-phenotype responses [54, 55, 56, 57, 58, 59, 60]. Existing multiscale models often represent EGFR activation through an intracellular ODE driven by the ligand concentration sampled at a representative cellular location, such as the cell center. This approximation provides a practical way to couple receptor signaling with cell-level behavior. However, incorporating EGFR-EGF binding kinetics requires the explicit representation of receptors, which we attempted in a previous work [61].

Here, we further refine this approach by calibrating the intracellular and molecular counts to available data and experiments. Then, we study the dynamics of the 3D spatially resolved model to derive a novel continuous framework that stratifies cells according to receptor occupancy. The framework is further reduced into a population dynamics model that describes the average number of active receptor, obtained using a moment reduction technique. The models are collectively applied to investigate the effects of perturbations in the EGFR-EGF interactions on tumor initiation, under a wide range of physiological conditions. Specifically, we quantify the minimal number of EGFR receptors that a cell needs to have to become cancerous, as a function of the cell phenotype, ligand-receptor affinity, and free EGF concentrations. The remainder of the paper is organized as follows: Section 2 introduces a 3D multiscale model as well as a new continuous model of tumor growth that explicitly integrate EGFR-EGF binding kinetics. It also provides implementation details such as parameterization, analysis as well as the numerical and programming methods. Section 3 presents the results of numerical simulations and analytical estimates, starting with the calibration of the models to experimental data and then analyzing the impacts of changes in the EGFR-EGF binding kinetics together on the extracellular and intracellular regulation of tumor initiation. The novelty, contributions, and limitations of this study are discussed in Section 4.

## 2 Methods and models

In this work, we introduce a hierarchy of multiscale models for cancer initiation with explicit receptor-ligand dynamics. The hierarchy links three levels of description: a 3D stochastic multicellular model, a receptor-structured continuous model, and a reduced population-level dynamical system. Each level serves a distinct role. The 3D model resolves spatial and stochastic ligand-receptor interactions at the cell surface. The receptor-structured model translates these microscopic interactions into population dynamics across receptor-activation states. The reduced model provides a lower-dimensional formulation for analytical threshold analysis.

Section 2.1 presents an extended version of a previously published 3D multicellular model that explicitly incorporates receptor domains and their spatial interactions with ligands [61]. We enhance this model by (i) calibrating molecular quantities to experimentally measured values and (ii) replacing the discrete intracellular regulation submodel with an experimentally-calibrated and computationally efficient alternative. Section 2.2 introduces a receptor-structured continuous model of tumor growth in which cells are stratified according to the number of active EGFR clusters. This formulation incorporates EGFR-EGF binding and unbinding as transitions between receptor-activation states. Each state is assigned a receptor-dependent division rate, allowing the continuous model to capture the saturation effects observed in the 3D multicellular simulations. Under simplifying assumptions, we then derive a reduced version of the receptor-structured model and analyze it to obtain analytical estimates for tumor-initiation thresholds. The relationships between the different modelling levels are summarized in Figure 2. Throughout this work, *N*_*r*_ denotes the physiological level of total EGFR receptor clusters per cell, while *E* denotes the extracellular EGF concentration used across the model hierarchy. In the 3D multiscale model, *N* represents the discrete number of active EGFR receptor clusters on an individual cell, whereas in the continuous receptor-structured and reduced models, *n* denotes the continuous active receptor-cluster state used to stratify the cell population.

**Fig 2.**
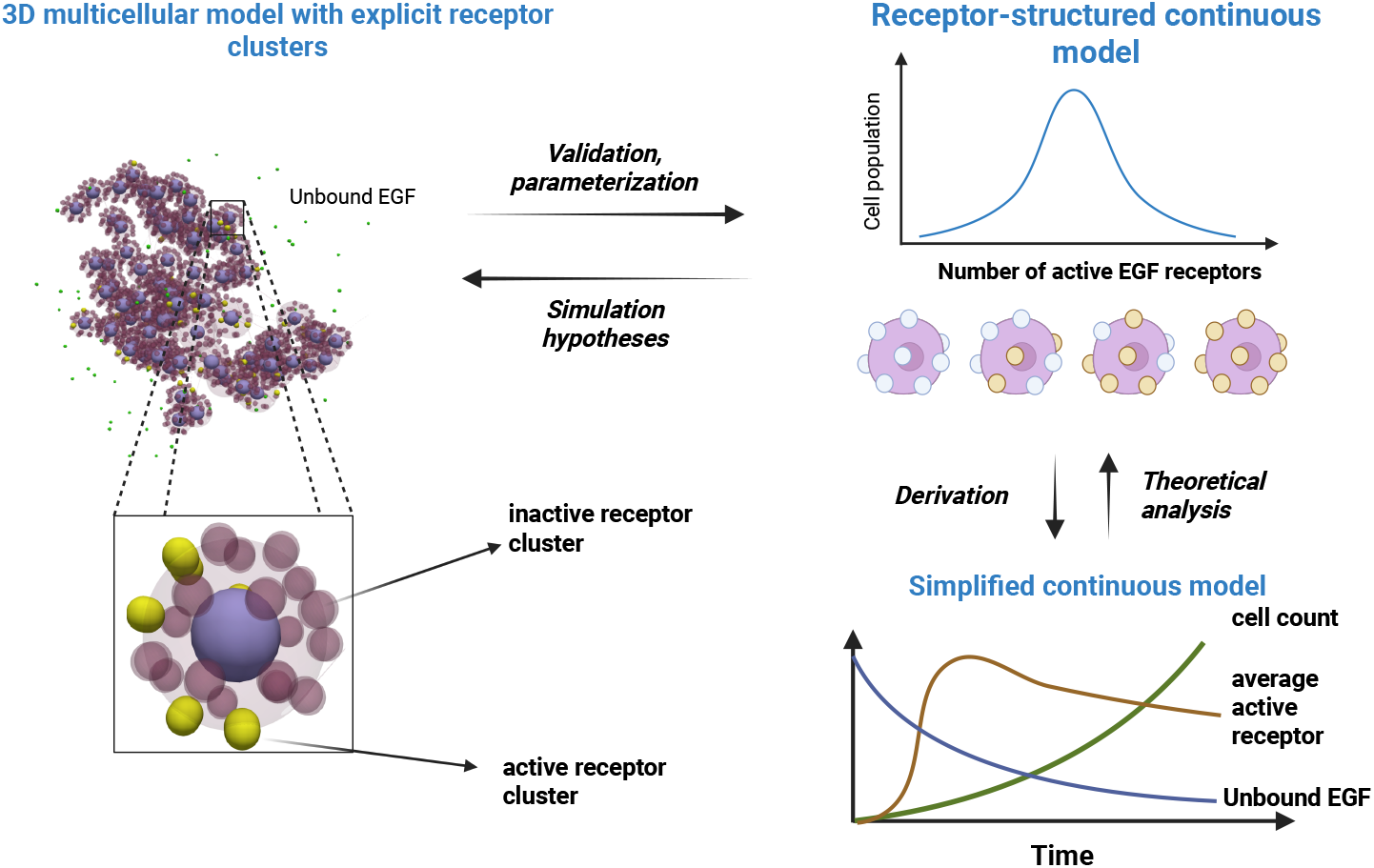
A summary of the interactions between the multiscale models used in this work to analyze the thresholds of tumorigenesis. A previous 3D multiscale model with explicit EGFR-EGF interactions is extended and calibrated to physiological EGFR and EGF ratios [61]. The model dynamics are analyzed to derive a continuous model structuring cells according to the number of active receptor. This model is used to further explore the dynamics of the system and test simulation hypotheses with the 3D multiscale model. The continuous receptor-structured framework is then reduced into a simplified ODE-based version, which is analyzed mathematically to produce analytical estimates for tumor initiation thresholds.

### 2.1 A 3D multicellular model of tumor growth with spatially-resolved EGFR-EGF interactions

In this study, we enhance a previous multiscale model that captures how mutational alterations in the EGFR receptors influence tumor initiation [61]. In the model, individual cells are represented as discrete spheres that can move, divide, and undergo apoptosis. Each cell membrane contains EGF receptors that bind to extracellular ligands (EGFs) and initiate the EGFR/ERK signaling cascade. At the intracellular level, we replace the previous discrete model with a computationally simple and experimentally calibrated ODE describing ERK activation. The terminal product, ERK, translocates to the nucleus, where it promotes the expression of transcription factors that govern cell fate decisions. Each simulations starts with one cell and takes place within a spherical computational domain with a radius of 50 *µm*. EGFs are also periodically introduced to the computational domain, and we calibrate their influx such that their numbers match experimental values. The model is refined by providing a more rigorous calibration for the physiological concentrations of EGFs and receptors.

#### 2.1.1 Extracellular regulation

We consider a spherical computational domain with a radius of 50 *µm*. EGF molecules are periodically introduced at random locations along the outer boundary of this domain. They are released at a constant rate such that, in the absence of cells, the total number of EGF corresponds to a physiological value. These proteins diffuse through the extracellular matrix and are removed from the domain upon degradation, with degradation times sampled from an exponential distribution. EGFs that reach the outer boundary are reflected into the domain. Each EGF and EGFR molecule is assigned a reaction radius, and binding occurs reversibly when these radii overlap. Upon binding, the EGFR becomes activated and initiates the cell’s intracellular signaling cascade.

In the model, each EGFR cluster is represented as two spheres anchored to the cell membrane. The axis connecting the centers of these spheres is oriented perpendicular to the tangent of the cell surface. Consequently, one sphere always remains outside the cell, while the other is positioned inside. Receptors are assumed to be uniformly distributed over the cell surface and can exist in two states: ‘on’ (active) and ‘off’ (inactive). An active receptor remains in the ‘off’ state for a duration corresponding to its residence time. The waiting times for EGF degradation and EGFR-ligand dissociation are sampled from exponential distributions, each characterized by its respective reaction rate. We refine the extracellular part of the model by incorporating realistic and physiologically relevant EGF and EGFR concentrations, such that the ratio of EGFR to EGF remains the same as detailed in B.

#### 2.1.2 Intracellular regulation

While activated, EGFR induces the EGFR/ERK cascade, which promotes the division of the cell. To speed up the computational cost, we replace the previous discrete implementation of the EGFR-ERK cascade with a simple continuous mode. The model tracks the number of active ERKs depending on the number of active EGFRs and natural decay, as follows:

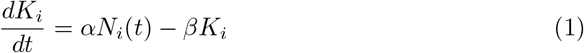

where the first term on the right-hand side of the equation describes the contribution of active EGFR clusters, with *N*_*i*_(*t*) denoting the number of the currently active EGFR clusters. The second term models the inactivation of ERK. The dynamics of ERK activation are calibrated using a lab experiment for transient EGFR activation [62] as shown in A. The cell divides if the number of active ERKs exceeds a certain threshold by the end of the G1-phase (*K*_*i*_ ≤ *K*\*).

#### 2.1.3 Cell movement and division

We adopt a center-based multicellular approach to describe tumor development [63]. Each cell is modeled as an elastic sphere containing an incompressible core representing its nucleus. The intracellular concentration of ERKs determines cell fate: if this concentration exceeds a defined threshold at the end of the G1 phase, the cell grows and divides by the end of its cycle; otherwise, it undergoes apoptosis. During division, the cell’s volume increases linearly until it doubles, after which two daughter cells are formed. The axis connecting their centers is assigned a random orientation to prevent spatial bias. The growth of the mother cell before division ensures that the daughter cells do not overlap with neighboring cells at the moment of their appearance. Each daughter cell inherits half of the mother cell’s intracellular proteins, while their coordinates are reinitialized randomly to reflect dilution effects at division. Tumor expansion arises from the mechanical interactions between cells during growth and proliferation. The duration of the cell cycle is set to 24 hours, with a small random perturbation uniformly sampled within the interval [− 3*h*, 3*h*]. The motion of individual cells is governed by Newton’s laws of dynamics:

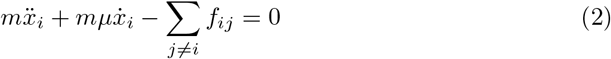

where *m* is the mass of the cell and *µ* is the friction constant due to the contact with the surrounding medium. The repulsive *f*_*ij*_ between two cells is computed using the following equation:

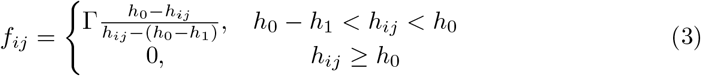

Here, *h*_*ij*_ is the distance between the centers of the two interacting cells i and j, *h*_0_ is the sum of their radii, *K* is a positive parameter, and *h*_1_ is the sum of the incompressible part of each cell. The force *f*_*ij*_ tends to infinity if *h*_*ij*_ decreases to *h*_0_ *− h*_1_. The same equation of Newton’s second law of dynamics and the force is used in the previous model. This cell-based model captures the biomechanics of tumor growth in environments characterized by varying Reynolds numbers. It can be simplified to a first-order formulation by assuming that the inertial term is negligible compared with the dissipative term [64, 50]. This approximation is particularly valid for tumors developing in microenvironments with low Reynolds numbers. In our model, however, we retain the inertial term to enable simulations of tumor growth across different anatomical locations with potentially distinct biomechanical conditions. We have previously used a similar approach to describe multiscale multicellular processes in 2D [65, 66, 67].

#### 2.1.4 Computer implementation

The code is implemented in C++ using the object-oriented programming paradigm. To ensure the accuracy of the Brownian dynamics implementation, we simulated the system with a decreasing time step until there is no significant change in mean behaviour [61]. The GNU Scientific Library (GSL) was used for the sampling of random variables. The intracellular ODE equation is solved using the Euler-explicit method.

The average CPU time of a numerical simulation corresponding to 15 days of tumour initiation is approximately 4 hours on an AMD Ryzen 7 machine with 36 Gb of internal memory, although it can vary significantly according to the number of cells. The visualizations were produced using the software ParaView. The source code for calibration simulations is publicly available at https://github.com/Romasa/Cancer3D_validation/tree/main. The parameters specific to the multicellular model are specified in Table 1.

**Table 1.** Values of the considered simulation parameters used in the simulations of the 3D multicellular model which were fixed in the current study.

| Parameter | Description | Value |
| --- | --- | --- |
| $\Delta t$ | Time step | 0.02 *min* |
| $r_{BD}$ | Reaction radius | 0.15 $\mu m$ |
| $d_{cell}$ | Cell radius | 4 $\mu m$ |
| $d_{nucleus}$ | Nucleus radius | 2 $\mu m$ |
| $\Gamma$ | Contact force coefficient | 1000 $kg \cdot \mu m^{-1} \cdot min^{-1}$ |
| $D_{EGF}$ | 360 | EGF diffusion coefficient [68] |
| $T_{cell}$ | 24 *h* + $\epsilon$ | Cell cycle duration |
| $\epsilon$ | Uniform from [-3h, 3h] | Cell cycle perturbation |

### 2.2 A continuous approach to tumor growth modelling with explicit EGFR-EGF interactions

We introduce, for the first time, a new modelling approach of cancer stratifying cells according to the number of active receptors. It allows the incorporation of receptor-ligand binding kinetics by assuming that the growth rate of cells depends on the number of active receptors that they express. The model is parameterized according to the molecular levels of the 3D multiscale model. It is reduced under some simplifying assumptions and analyzed to derive estimates for tumor growth initiation.

#### 2.2.1 A receptor-structured model of tumor initiation and progression

We introduce a continuous model that captures ligand-receptor dynamics and their impact on tumor growth. This model extends classical tumor growth frameworks by incorporating ligand-receptor interactions as transitions between different active receptor states. Tumor cells are structured according to the number of active EGFR clusters they harbor. We define the space [0, *N*_*r*_] to represent the number of active EGFRs, where *n* denotes the number of active receptor clusters, and *N*_*r*_ represents the total number of receptor clusters per cell. The model explicitly tracks the concentration of free EGFs and describes how EGF-EGFR binding and unbinding events drive cell redistribution across the space [0, *N*_*r*_]. Specifically, cells with fewer active receptors exhibit higher EGF-binding rates, accelerating their transition to states with more active EGFRs. Simultaneously, all cells transition to states with fewer phosphorylated EGFRs at a rate that depends on the dissociation constant and the number of active clusters. The growth rate of tumor cells is governed by the number of active receptors and the availability of growth factors. We describe the tissue-level dynamics of free EGFs as follows:

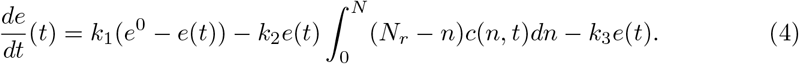

Here, the first term on the right-hand side of the equation describes the influx of EGF molecules to the tissue. The second represents the consumption of EGFs by free EGF receptors. It depends on the number of cells and their corresponding free receptors. The third term describes the natural degradation of EGF molecules. The initial concentration will be set to the physiological range (*e*^0^). Next, we will describe the concentration of cancer cells, stratified according to the number of active EGFR molecules (*n*):

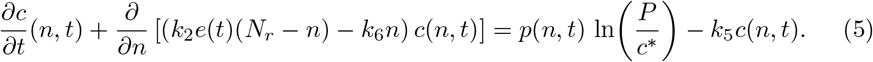

Here, the second term on the left-hand side describes the transition of cells to states with a higher number of phosphorylated EGFR receptors, driven by the binding of free EGFRs to available EGF molecules, and their movement towards states with fewer active EGFRs, occurring at a speed equal to the dissociation constant multiplied by the number of active receptors. The first term on the right-hand side accounts for the proliferation of cancer cells, which depends on the number of active receptors they carry (*n*). This proliferation considers a Gomperzian growth, with *P* and *c*\* being the population capacity and total population of cells, respectively. It considers that daughter cells inherit half of the active receptors of the mother cells following mitosis:

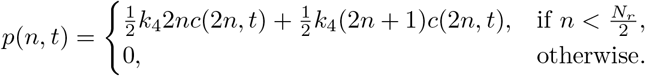

Here, *k*_4_ represents the cell growth rate per active receptor cluster and can describe the genotype-driven aggressiveness of the cell. The last term on the right-hand side of Equation (5) represents the apoptosis of cancer cells. We prescribe zero-flux boundary conditions and consider a uniform distribution of cells from *n* = 0 to *n* = 1 with a total density equal to one.

#### 2.2.2 Model reduction

Under the assumptions that (i) tumor growth is exponential and its rate correlates linearly with the number of active receptors per cell, and (ii) daughter cells inherit the same number of active receptors as the mother cell, the system (4),(5) dimensionality can be downsized using moment-reduction [69, 70]. Considering the mean number of active receptor clusters as the first moment, we obtain the following ODE system:

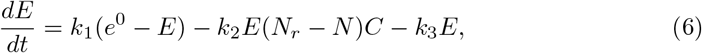

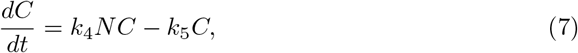

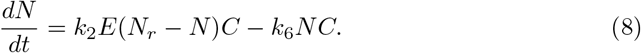

Here, *E* is the concentration of EGF, *C* is the number of cancer cells, and *N* represents the average number of active receptors per cell. The reduced system tracks the results of the receptor-structured model, as shown in Figure 3.

**Fig 3.**
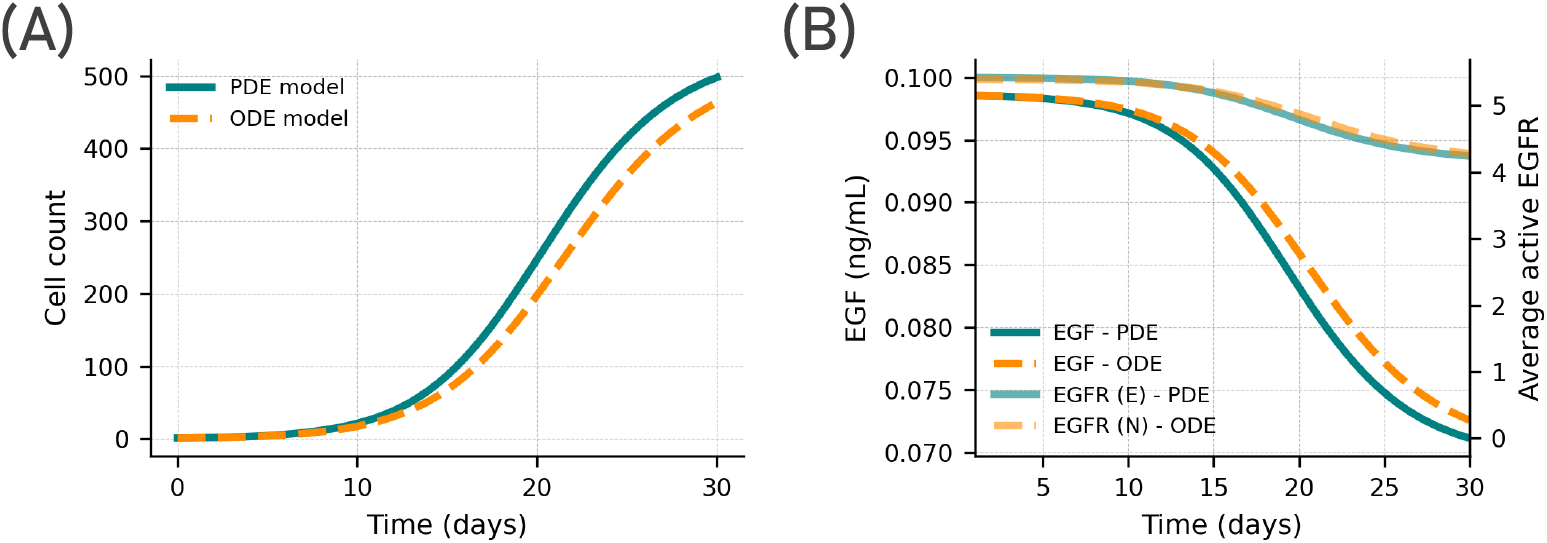
A comparison between the results obtained using the PDE-ODE model and the reduced ODE system.

#### 2.2.3 Analysis of the reduced model

We calculate the steady-state equilibrium points of the system (6)-(8) by setting:

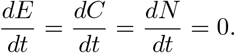

The system has three equilibrium points, given by the following solutions:

- Equilibrium (1): 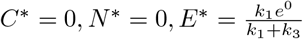
- Equilibrium (2): 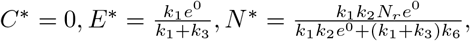
- Equilibrium (3): 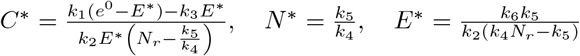

The first equilibrium points correspond to tumor-free scenarios. The third represents a tumor growth case. The stability analysis given in Appendix C shows that the first equilibrium is stable, the second is conditionally stable, and the third is stable under physiological conditions. This equilibrium is only physiologically possible if *E*\* > 0 and *C*\* > 0. Therefore, it is only possible if the following conditions on the initial EGFR and EGF concentrations are met:

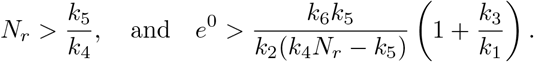

These conditions show that the minimum number of EGFR clusters required for tumor initiation depends solely on the intrinsic fitness of the cells: more aggressive cells need fewer receptors to trigger tumorigenesis. At the same time, the amount of available EGF required to initiate growth decreases as the number of EGFR clusters increases, and it decreases further when the EGFR–EGF binding affinity becomes stronger.

A single summary quantity that can predict whether a tumor will grow or regress is the basic reproduction number. Although this concept was initially developed for infectious disease modelling, it has also proven useful for characterizing tumor growth dynamics [71, 72]. We estimate the basic reproduction number (*R*_0_) by computing the average number of new cancer cells produced by a single cell at the earliest stage of growth, when EGF is abundant (*E* ≈ *e*^0^). Under this assumption, the number of new cells generated per cancer cell at the onset is:

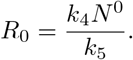

Here, *N* ^0^ represents the number of active receptors at the beginning. It can be calculated by taking *E* = *e*^0^ in Equation (8) at steady-state:

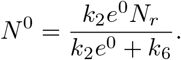

Thus, the basic reproduction number:

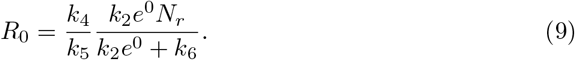

It indicates that tumor growth occurs only when the threshold exceeds 1, thereby defining the conditions under which proliferation is sustained as a function of the intrinsic division potential, the EGFR–EGF unbinding kinetics, and the local availability of EGF.

#### 2.2.4 Numerical implementation

The receptor-structured model was implemented using the finite difference method, with a time and a space steps equal to *dt* = 0.005 hour and *dx* = 0.25 receptor cluster, respectively. The explicit Euler scheme was used for time discretization. The transport terms were discretized using an upwind scheme. The reduced ODE system was implemented using the Euler explicit method. The code was implemented in Python using the Jupyter notebook.

## 3 Results

### 3.1 Calibration of the models to experimental data

We begin by calibrating the receptor-structured models against experimental results obtained from an *in vivo* tumor system in nude mice [73]. In that study, the efficacy of the chemotherapeutic agent CPT-11 was assessed in tumors with and without EGFR overexpression. We used the vehicle-treated group to calibrate the parameters of our two modeling frameworks. Specifically, we first initialized all parameters using values reported in the literature, and then tuned the cellular aggressiveness to reproduce the baseline growth curve. This calibration was achieved by adjusting the threshold of ERK molecules required for cell division (*K*\*) in the hybrid model and modifying the division rate parameter (*k*_4_) in the receptor-structured model. After that, we increased the number of receptors in each model to match the experimentally observed growth rates during EGFR overexpression. The numerical values of calibrated parameters for the 3D multicellular and receptor-structured models are provided in Tables 2 and 3.

**Table 2.** Parameters used to validate the results of the 3D multicellular model.

| Parameter | Value (baseline) | Value (EGFR overexpression) | Description |
| --- | --- | --- | --- |
| $N_r$ | 6 | 24 | EGFR clusters per cell |
| $\alpha$ | $2285.3 \text{ cluster}^{-1} h^{-1}$ | $2285.3 \text{ cluster}^{-1} h^{-1}$ | ERK activation rate by EGFRs (fitted to [74]) |
| $\beta$ | $6.251 h^{-1}$ | $6.251 h^{-1}$ | ERK degradation rate (fitted to [74]) |
| $K^*$ | 50000 | 50000 | ERK threshold enabling cell division |
| $d_{EGF}$ | $0.69 h^{-1}$ | $0.69 h^{-1}$ | EGF degradation [68] |
| $d_{EGFR}$ | $0.42 h^{-1}$ | $0.42 h^{-1}$ | EGFR-EGF unbinding constant [75] |

**Table 3.** Parameters used to validate the results of the receptor-structured continuous model. Values were derived directly from the 3D multiscale model simulations or through fitting to its results.

| Parameter | Value (baseline) | Value (EGFR overexpression) | Description |
| --- | --- | --- | --- |
| $e^0$ | 197 | 197 | EGF molecular count |
| $N_r$ | 6 | 24 | Number of EGFR clusters |
| $k_1$ | $50 \text{ ng.ml}^{-1}.h^{-1}$ | $50 \text{ ng.ml}^{-1}.h^{-1}$ | EGF influx rate |
| $k_2$ | $0.0025 \text{ ml.ng}^{-1}.h^{-1}$ | $0.0025 \text{ ml.ng}^{-1}.h^{-1}$ | EGF-EGFR binding rate |
| $k_3$ | $0.69 \text{ h}^{-1}$ | $0.69 \text{ h}^{-1}$ | Rate of EGF degradation |
| $k_4$ | $0.0145 \text{ h}^{-1}$ | $0.0157 \text{ h}^{-1}$ | Rate of cell division |
| $k_5$ | $1/24 \text{ h}^{-1}$ | $1/24 \text{ h}^{-1}$ | Apoptosis rate |
| $k_6$ | $1/1.5 \text{ h}^{-1}$ | $1/1.5 \text{ h}^{-1}$ | EGFR inactivation rate |

After their calibrations, the models produce growth patterns that are consistent with the experimental tumor growth rates for cells with 6 receptor clusters (baseline) and 24 clusters (EGFR overexpression), as shown in Figure 4. The receptor-structured model provides a closer match to the experimental trajectories than the 3D multicellular model, largely due to its inclusion of Gompertzian growth dynamics. Analysis of the receptor-structured simulations showed that cells preferentially accumulate around 2.5 receptor clusters in the baseline case and around 10 receptor clusters under EGFR overexpression. The 3D multicellular predictions represent the median and 95% confidence intervals computed from 25 independent simulations.

**Fig 4.**
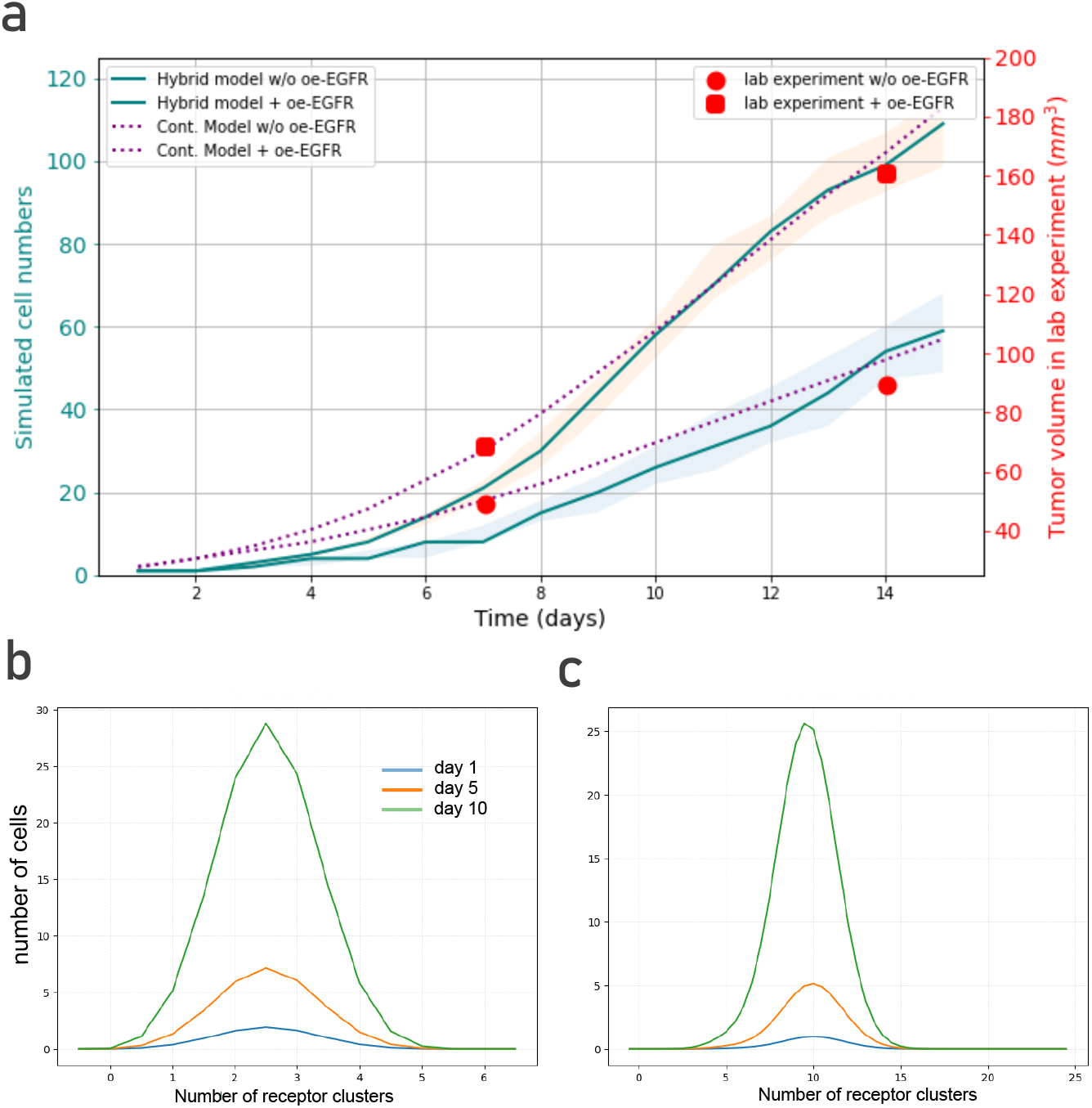
(a) Calibration of the developed models against *in vivo* experiments of xenografted tumor growth in nude mice, with and without EGFR overexpression. Experimental tumor volumes (red points) are reproduced from Figure 5 of Yin et al. (2023) [73]. Model predictions for wild-type cells and EGFR-overexpressing cells correspond to simulations using 6 and 24 receptor clusters, respectively. The 3D hybrid simulation results display the median tumor cell count together with the 95% confidence interval as ribbons. (b–c) Results from the receptor-structured model showing the distribution of cells across receptor-number classes at three representative time points, in the absence (b) and presence (c) of EGFR overexpression.

We analyzed the 3D multicellular simulations to gain insights into the spatial effects of tumor growth (Figure 5). EGFR overexpression was modeled by increasing the number of receptor clusters per cell from 6 (Figure 5-a) to 24 Figure 5-d). Without overexpression, the tumor grows from 14 cells on day 5 (Figure 5-b) to 22 cells on day 10 (Figure 5-c). A spherical morphology emerges, with fewer cells in the center due to limited EGFR availability. With EGFR overexpression, the tumor reaches 19 cells on day 5 (Figure 5-e) and 60 cells on day 10 (Figure 5-f). These simulations also show a spherical structure, with denser growth at the periphery. The cell-number ratio between the two conditions increases over time. This highlights the accelerating effect of EGFR overexpression on tumor expansion, which can also be observed experimentally.

**Fig 5.**
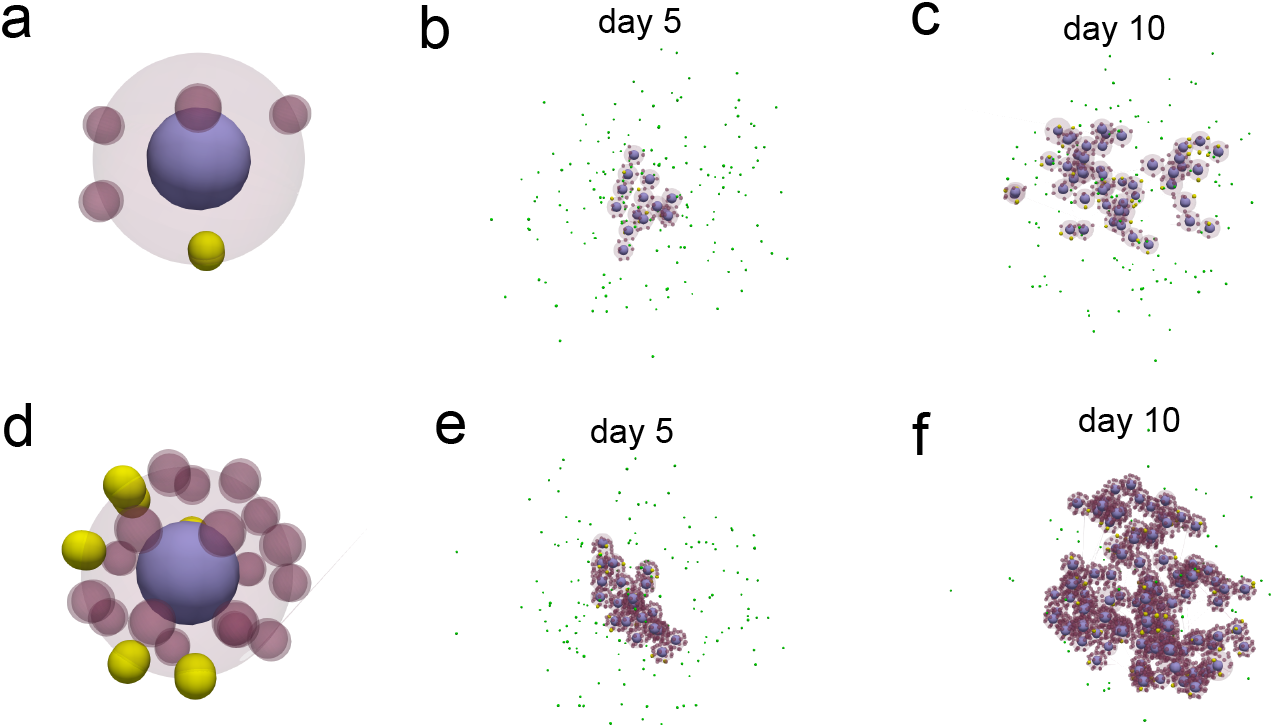
(a-d) Snapshots of individual cells used in the numerical simulations for the baseline (a) and EGFR overexpression (b) conditions. Cells contain 6 and 24 receptor clusters, respectively, with active receptors shown in solid yellow. (b-c) Screenshot of the validation simulations with the 3D multicellular model for the baseline case on days 5 and 10. (e-f) Results of validation simulations conducted for the case of EGFR overexpression on days 5 and 10.

### 3.2 A reduced EGF concentration is required for tumor initiation during EGFR overexpression

After validating both frameworks, we used them to identify the EGF concentrations that permit tumor growth under different receptor-cluster configurations. Lower EGF concentrations are typically observed in patients with chronic medical conditions such as kidney disease [76], metabolic disease [77], and advanced age [78]. Higher EGF and EGFR levels increase the likelihood of ligand-receptor binding and thus promote cell division. To determine the growth threshold in the 3D multicellular model, we tested four physiologically low EGF concentrations (0.05-0.2 ng/ml) and analyzed the number of cells on day 15 across 25 simulations per condition (Figure 6, a-d). For each simulation set, we calculated the 25%, 50%, 75% quartiles and outliers. At 0.05, ng/ml, the tumor went extinct in all simulations, regardless of receptor-cluster number. At 0.1 ng/ml, tumors grew only when cells had at least 16 clusters. Increasing the EGF concentration lowered this requirement: tumors initiated with 9 clusters at 0.15 ng/ml and with 6 clusters at 0.2 ng/ml. The impact of increasing EGFR cluster number diminishes when EGF availability is limited, as illustrated in Figure 6-c and d.

**Fig 6.**
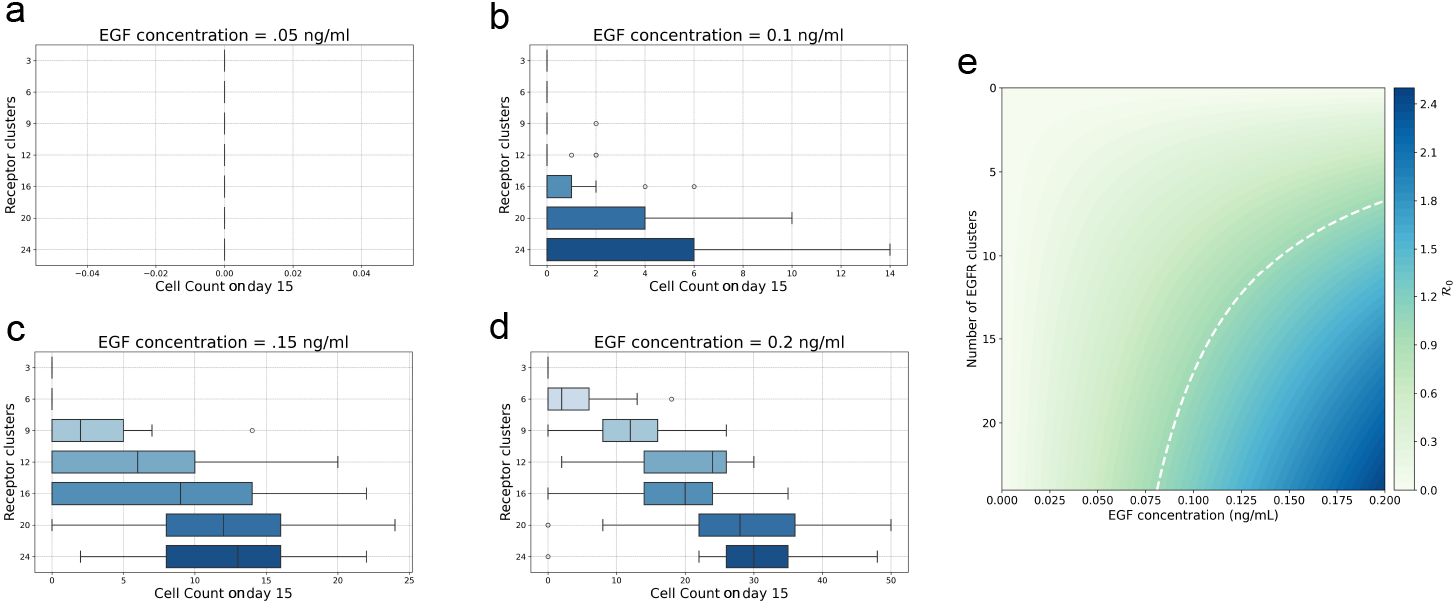
(a-d) Cell counts on day 15 across 25 simulations for different EGF concentrations and EGFR cluster numbers. Boxplots represent variations across simulations, including the quartiles, median, and outliers. (e) Basic reproduction numbers computed using the simplified continuous model for the corresponding EGF and EGFR molecular levels. The white dashed line indicates the border between regions where *R*_0_ is greater or less than one.

Next, we analyzed tumor initiation in the receptor-structured model using basic reproduction number estimates from the reduced form. The results in Figure 6-e show that for cells with fewer than 24 receptor clusters, EGF concentrations below 0.075 ng/ml typically lead to tumor extinction. Increasing the EGF concentration reduces the number of receptors required for tumor initiation, closely matching the trends observed in the 3D multicellular model. Specifically, EGF levels of 0.1, 0.15, and 0.2 ng/ml require roughly 18, 10, and 7 receptor clusters, respectively. These findings also confirm that the marginal impact of additional EGFR clusters decreases when EGF is abundant, consistent with the 3D simulations.

### 3.3 Constitutive EGFR activation reduces the EGF and EGFR count necessary for tumor initiation

Several EGFR mutations found in cancers can further shift this balance by promoting constitutive receptor activation. These alterations often stabilize the active conformation of the receptor or enhance spontaneous dimerization, allowing signaling to proceed for a long time following an initial activation. As a result, mutant EGFR variants can display higher affinity to EGF, making them potent drivers of uncontrolled growth [79]. In addition to mutations, EGFR clustering also promotes the sensitivity to EGF molecules and increases EGFR-EGF affinity [80].

To assess the role of EGFR-EGF affinity in tumor initiation, we analyzed numerical simulations from both the 3D multicellular and receptor-structured models. In the 3D model, we varied the EGF-EGFR unbinding constant so that the complex remained bound for periods ranging from 12 hours (*d*_EGFR_ = 0.00139 min^−1^) to 100 minutes (*d*_EGFR_ = 0.01 min^−1^). We performed 25 simulations per condition and kept all other physiological parameters identical to the validation case, including an EGF concentration of 0.4 ng/ml. For each unbinding constant, we examined both the baseline (6 clusters) and EGFR overexpression (24 clusters) scenarios. Figure 7-a shows a steep decline in the number of cells on day 15 as the unbinding constant increases. At the highest value (*d*_EGFR_ = 0.01 min^−1^), tumor growth is fully suppressed in both receptor configurations. At lower unbinding rates, tumors initiate and reach larger final sizes as the EGF-EGFR binding time increases.

**Fig 7.**
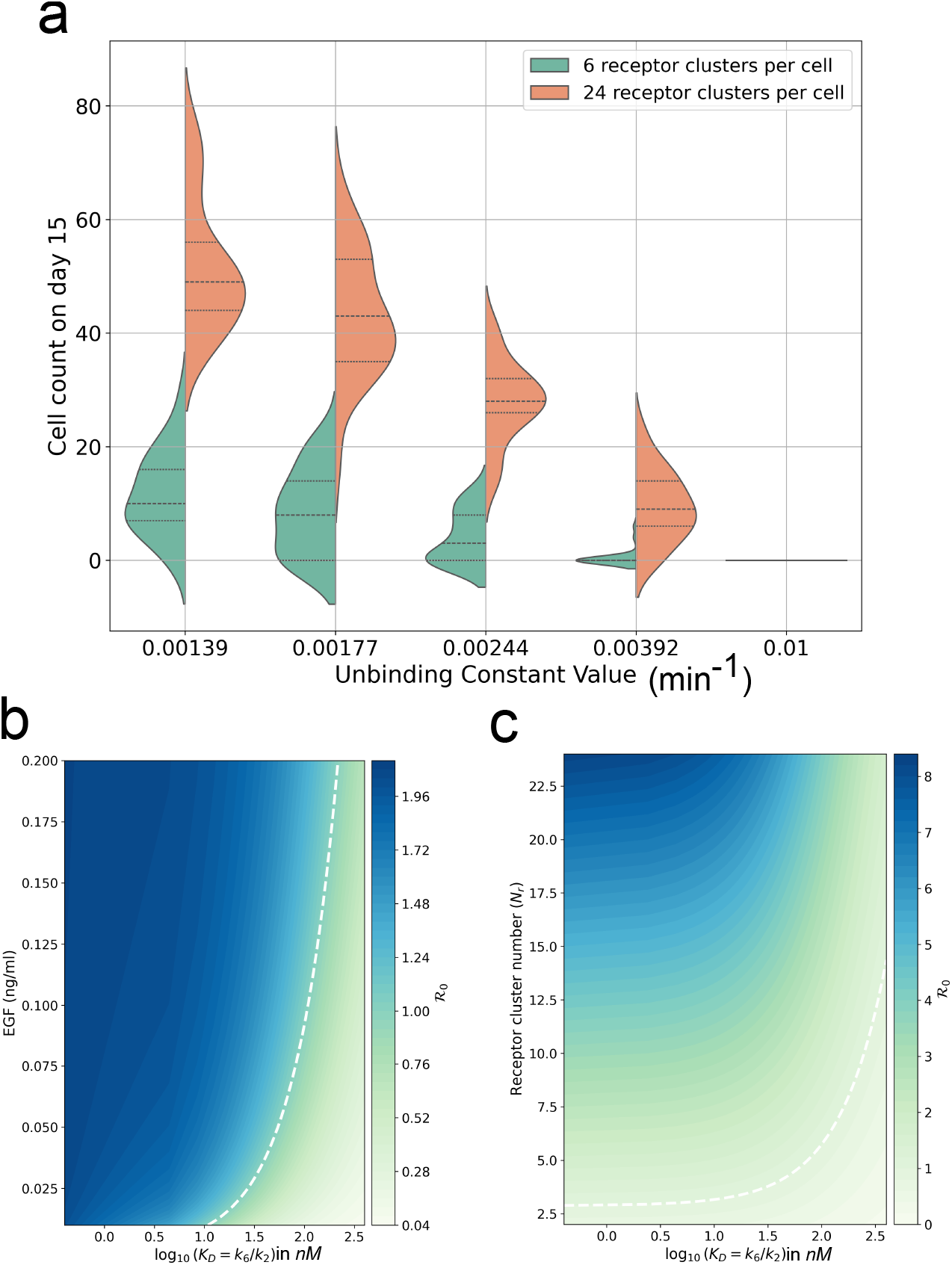
(a) A summary of the 3D model simulation results showing the distribution of cell numbers on day 15 across 25 simulations in the baseline and EGFR overexpression cases when varying the EGF-EGFR unbinding constant. (b) Basic reproduction number estimates from the reduced form of the receptor-structured model representing the chances of tumor growth in a cell with 6 receptor clusters as a function of the EGF concentration and the EGFR-EGF dissociation constant. The white dashed line indicates the border between regions where ℛ_0_ is greater or less than one. (c) Basic reproduction number estimates as a function of the number of EGFR clusters and the dissociation constants when considering a constant EGF concentration of 0.1 ng/ml.

We used the basic reproduction number of the receptor-structured model to examine tumor initiation across a broader range of physiological conditions. Specifically, we varied the equilibrium dissociation constant (*K*_*D*_), defined as the ratio of the unbinding and binding rates, along with the EGF concentration and the number of EGFR clusters (Figure 7-b,c). In all cases, increasing *K*_*D*_ reduced the likelihood of tumor initiation, with the threshold depending on both EGF availability and receptor abundance. For all tested EGF concentrations, tumor growth required *K*_*D*_ ≥ 1, nM, and this threshold increased as EGF levels rose. Tumor initiation was also prevented when the receptor number was minimal (below 2.5 clusters per cell), even under high-affinity binding. Increasing the number of receptor clusters promoted tumor initiation and raised the maximal *K*_*D*_ compatible with growth. These results show that the dissociation threshold depends jointly on extracellular ligand availability and receptor density.

### 3.4 Aggressive cell phenotypes require fewer receptors to initiate tumorigenesis

Finally, we examined how cell aggressiveness, defined by its intrinsic division potential, shapes the thresholds for tumor initiation. Genetic alterations that prolong the duration and strength of intracellular signaling drive this aggressiveness. In the 3D multicellular model, such alterations could correspond to mutations in K-Ras, N-Ras, or B-Raf oncogenes, which amplify ERK activation following EGFR stimulation. To study this effect, we performed numerical simulations across nine values of the ERK activation rate by EGFR receptors (*α*). We performed 25 simulations for each ERK activation value, and retained the parameters used in the validation scenario, with an EGF concentration of 0.4 ng/ml and six receptor clusters per cell. Figure 8-a shows that increasing *α* beyond a threshold of approximately 5,000 cluster^−1^min^−1^ determines whether cells ultimately undergo extinction or initiate tumor growth. We then assessed how this threshold shifts with greater extracellular ligand availability by increasing the EGF concentration. Figure 8-b illustrates that higher EGF levels promote tumor growth even at lower values of *α*. These findings demonstrate that the ERK-activation threshold enabling tumor initiation depends on the extracellular supply of ligands.

**Fig 8.**
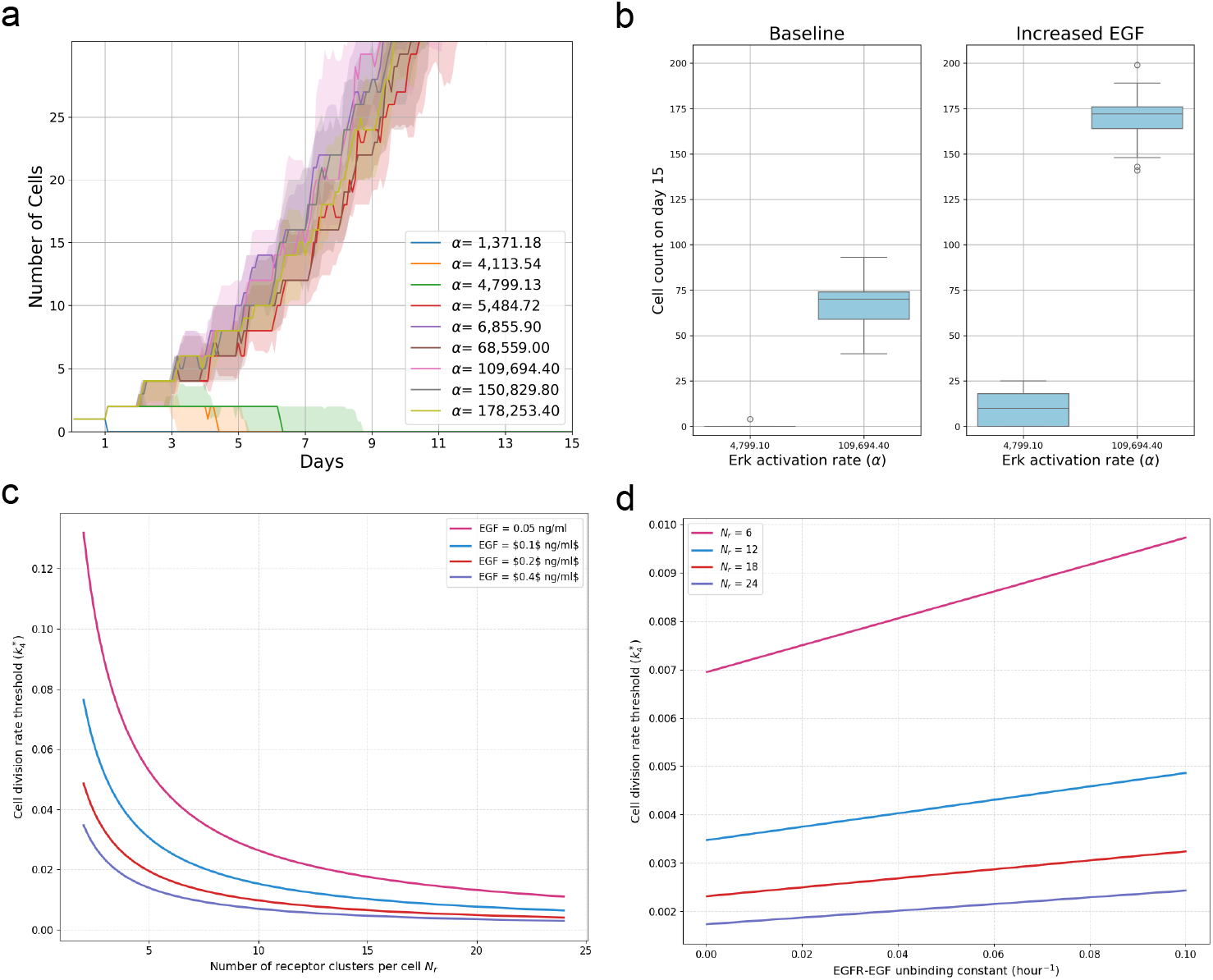
(a) Results of numerical simulations with the 3D multicellular model showing the number of cells over time for different ERK activation values (*α*). (b) The number of cells on day 15 in numerical simulations for baseline (0.4 ng/ml) and increased (0.6 ng/ml) EGF concentrations.

To further examine the effects of cell aggressiveness, we used the basic reproduction number approximation (Equation 9) to identify the threshold division rate per receptor cluster, *k*_4_, required for ℛ_0_ > 1:

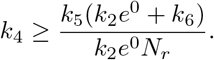

We then evaluated how this threshold varies with key biophysical parameters. Figure 8-c shows that increasing the EGF concentration lowers the threshold division rate, indicating that cells require weaker proliferative signals to initiate growth under ligand-rich conditions. This trend holds across cells with different numbers of receptor clusters, with highly clustered cells exhibiting an even lower threshold. We also quantified how this threshold depends on the unbinding constant. The reduced model predicts a positive linear relationship between the threshold division rate and the unbinding constant, as shown in Figure 8-d.

## 4 Discussion

Existing discrete and continuous tumor growth models often simplify receptor dynamics by embedding them into aggregate parameters, such as growth rates or effective counts of active receptors, to reduce complexity and computational cost. This paper introduces complementary modeling frameworks that explicitly capture EGFR-EGF interactions.

The contribution of these frameworks is that they explicitly link extracellular ligand availability, receptor abundance, and binding affinity to cellular proliferative potential, which enables the exploration of the ligand-receptor mechanisms driving tumor initiation. Together, the frameworks provide numerical and analytical estimates of the thresholds required for tumor initiation, depending on the balance between pro- and anti-proliferative factors governed by EGFR and EGF levels, EGFR-EGF affinity, and intracellular regulatory capacity.

The first framework is a 3D multicellular model in which each cell includes membrane domains representing EGFR receptor clusters. EGF ligands are represented as discrete particles that move by Brownian dynamics and interact with cell-bound EGFR clusters. As a result, receptor activation emerges from spatially localized and stochastic ligand-receptor interactions at the cell surface. Many hybrid multiscale models embed receptor activation inside intracellular ODEs driven by ligand concentrations sampled at a single point, usually the cell center [81, 82, 83]. Our approach complements these formulations by explicitly resolving receptor-ligand encounters in three-dimensional space. This assumption enables the incorporation of the role of the exposed area of the cell and the number of receptors it contains in driving the strength of the signaling. Although this model was previously introduced [61], we improve it here by replacing the discrete intracellular regulation module with a simplified and experimentally validated ODE system. A key limitation remains: true molecular counts of EGFR and EGF are out of reach because agent-based simulations are computationally expensive. To manage this, we applied coarse graining and calibrated the effective molecular numbers to preserve physiological EGFR-to-EGF ratios. This allowed us to reproduce the essential dynamics of EGFR and its overexpression while keeping the model computationally feasible. Although the framework is computationally intensive, it gave us direct insight and quantification of the spatial and stochastic effects that shape the earliest stages of tumor growth.

The second framework is a receptor-structured continuous model, developed here for the first time. Previous mathematical oncology studies have used structured population models, typically cell-cycle [84], size-[85], molecular expression- [86], or phenotype-structured PDEs [87, 88], to represent intratumoral heterogeneity and link intracellular regulation to population-scale behavior. We adopt this approach by structuring the cell population according to receptor occupancy, which is a key determinant of signaling activation (e.g., EGFR/ERK) and cell division. Our structured framework links cell phenotype, EGF-EGFR affinity, and the population-scale ligand availability within a unified and spatially-homogeneous model. We also derived a reduced form using moment-based simplifications. This reduced model reproduces the dynamics of the full receptor-structured model and allows mathematical analysis of the conditions required for tumor initiation. The two continuous models are readily extensible and can incorporate additional tumor-growth features, including alternative growth laws, cell dormancy, immune interactions, and treatment effects as well as additional receptor states such as blocked or decoy.

Figure 9 compares the tumor-initiation thresholds predicted by the reduced ODE model and the full 3D stochastic multicellular model. The reduced model captures the inverse relationship between extracellular EGF availability and the number of EGFRs per cell required for tumor initiation. The thresholds obtained from the 3D simulations follow the same qualitative trend, indicating that increasing EGF availability lowers the receptor density needed to sustain tumor growth. A moderate discrepancy is observed at higher EGF concentrations, where the reduced ODE model predicts slightly lower threshold values than the 3D simulations. This difference may reflect the distinct assumptions and resolution of the two model formulations. In the 3D stochastic model, spatial heterogeneity allows cells near the tumor boundary to encounter and bind ligands differently from cells in the tumor interior, which can affect the distribution of receptor activation across the population. In contrast, the reduced ODE model represents the population through averaged variables and assumes an effective growth law, which does not explicitly resolve spatial ligand gradients, local receptor-ligand encounters, or variability in receptor activation during cell division. Therefore, the discrepancy likely reflects the expected loss of spatial and growth assumptions during model reduction, rather than a qualitative disagreement between the two approaches.

**Fig 9.**
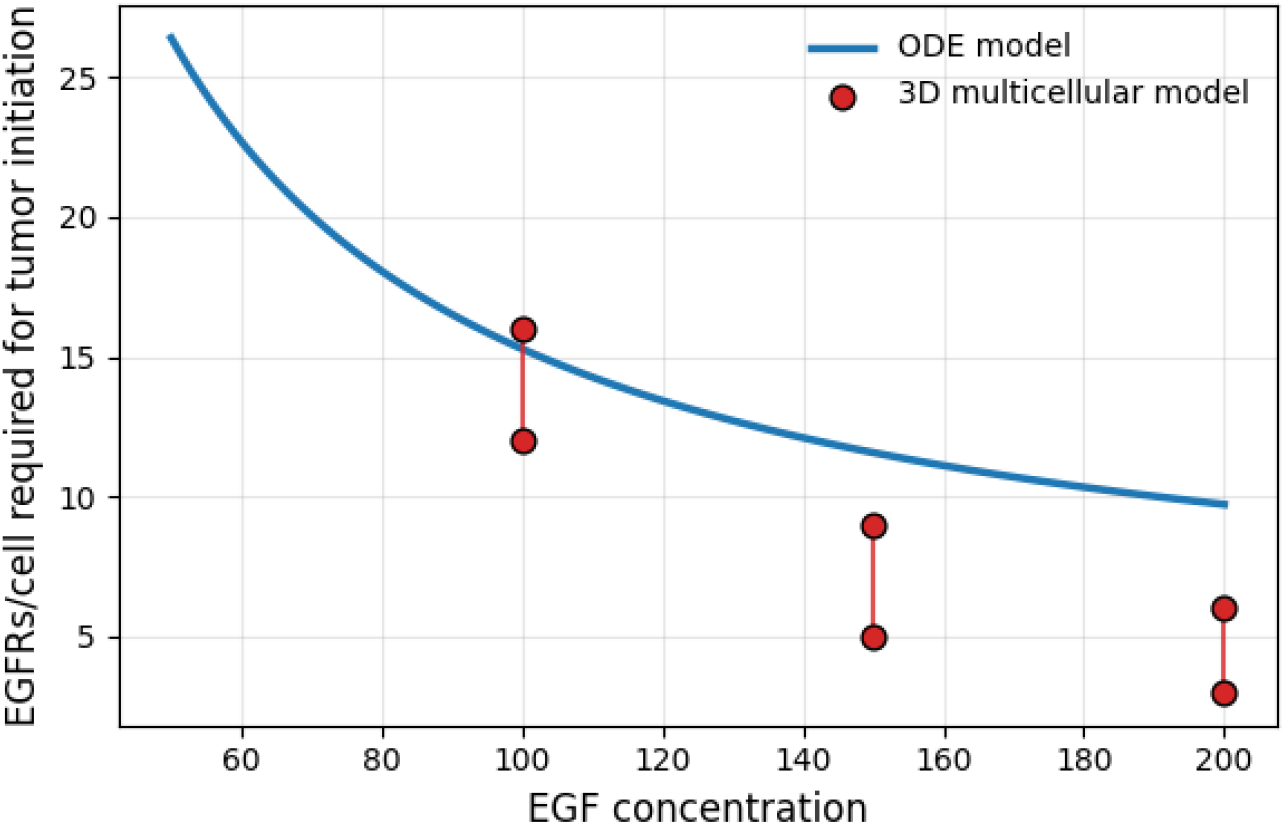
The number of EGFR clusters per cell required for tumor initiation under different EGF concentrations according to the 3D multicellular and reduced ODE models.

After validating the models against *in vivo* tumor growth data with and without EGFR overexpression, we used both frameworks to estimate the EGF and EGFR molecular counts required for tumor initiation. As expected, cells with higher receptor numbers required lower EGF concentrations to initiate growth because they are able to bind more ligands. The models also showed that the marginal contribution of additional receptors decreases at EGF abundance, consistent with the negative cooperativity reported for EGFR [89]. Further, our predictions also agree with experiments showing that the abundance of EGFR ligands can be a driver of tumor growth [90, 91]. Notably, both the 3D multicellular and receptor-structured models produced similar thresholds for both EGF and EGFR, suggesting that the receptor-structured model and its reduced version can serve as a surrogate for the more detailed and computationally intensive 3D multicellular model.

We next examined the role of EGFR-EGF affinity in tumor initiation. Although previous studies have shown that activating EGFR mutations (e.g., L858R, exon 19 deletions) enhance proliferation and tumorigenesis [92], the quantitative impact of affinity on tumor initiation across different EGFR and EGF levels remains unclear. Our models revealed that higher affinity reduces the molecular EGF and EGFR counts required for tumor initiation. Finally, our models indicate that cells require weaker proliferative signals to initiate growth under EGFR overexpression, ligand-rich conditions, or constitutive EGFR-EGF constitutive binding. This condition corresponds to the cases of aberrant EGFR-ERK signalling, which drives tumor growth in the absence of EGFR overexpression or EGF abundance [93].

Table 4 highlights the complementary roles of the three modelling frameworks. The 3D multiscale model provides the highest level of mechanistic fidelity, explicitly incorporating stochastic receptor–ligand interactions, spatial effects, and cell heterogeneity. It is therefore most appropriate when integrating data from multiple biological scales or when investigating the combined influence of molecular variability, spatial organization, and phenotypic diversity. However, its computational cost limits extensive parameter exploration and theoretical analysis.

**Table 4.** Comparison of the strengths and limitations of the proposed modeling frameworks.

|  | 3D Multiscale | Receptor-structured | Reduced ODE |
| --- | --- | --- | --- |
| Accuracy | High | High | Moderate |
| CPU Time | hours | minutes | milliseconds |
| Theoretical Analysis | Limited | Partial | Extensive |
| Stochastic | Yes | No | No |
| Spatial | Yes | No | No |
| Heterogeneity | Yes | Partial | No |

The receptor-structured continuum model offers a compromise between biological detail and tractability. By structuring cells according to receptor occupancy, it captures heterogeneity in activation states while eliminating explicit spatial and stochastic components. This formulation is particularly useful for analyzing the distribution of receptor activation across the population and for studying how heterogeneity influences growth thresholds at moderate computational expense. The reduced population dynamics model provides the most computationally efficient representation. Although it neglects stochastic effects, spatial structure, and explicit heterogeneity, it preserves the essential coupling between receptor activation and proliferation through the average number of active receptors. This low-dimensional system enables rigorous theoretical analysis, including stability and threshold characterization, and facilitates rapid parameter exploration.

It is important to note several limitations of the models introduced here. In the 3D multicellular model, we imposed simplifying assumptions to enable simulations of early tumor growth. Although small tumors with few cells can be modeled accurately, large-scale simulations are computationally prohibitive on standard hardware. We therefore reduced the number of EGFR and EGF molecules to lower computational cost using a coarse graining approach. The model also assumes that receptors consist of two clamped spheres, although a cylindrical geometry would be more realistic. This assumption was made because the spherical geometry makes it easier to computer reaction areas. Further, the current formulations do not explicitly account for receptor trafficking, internalization, or recycling dynamics. Instead, receptor concentration is assumed to remain in a quasi–steady state over the time scale of tumor initiation. Similarly, the downstream EGFR–ERK signaling pathway is represented in a simplified manner, and detailed intracellular regulatory cascades are not explicitly resolved. These processes are known to influence receptor availability and signal propagation and may alter quantitative threshold predictions. However, their inclusion would substantially increase model complexity and introduce additional poorly constrained parameters. The present study focuses specifically on isolating the dynamical contribution of receptor–ligand occupancy to tumor initiation. Finally, the model assumes that receptor cluster can be either “on” or “off”, while in reality they can be at partial activation states. In the future, we will extend the model by describing the activation state of each cluster using an ODE, which depends on local EGF concentrations. In the receptor-structured model, we did not account for saturation effects when many receptors become activated. This approximation was necessary to remain consistent with the intracellular model embedded in the 3D framework.

## 5 Conclusion

The proposed models are used to distangle the effects of EGF availability, EGFR abundance, receptor-ligand affinity, and cellular proliferative potential on tumor initiation. Rather than identifying a single dominant mechanism, the models suggest that tumor onset may depend on coordinated effects across extracellular growth-factor availability, receptor activation, and cell phenotype. In this sense, the frameworks collectively provide a quantitative tool for exploring how changes in ligand-receptor interactions may shift tumor-initiation thresholds. The models also provide a basis for estimating how partial receptor blockade may influence tumor persistence or elimination under different biological conditions. In particular, the predicted level of EGFR inhibition required to suppress growth depends on cellular division potential, EGF availability, and EGF-EGFR binding affinity. These predictions should be interpreted as mechanistic hypotheses that require further experimental and clinical validation, but they may help identify parameter regimes in which anti-EGFR strategies are expected to be more or less effective.

Future extensions could integrate the proposed framework with patient-specific measurements, including growth-factor levels, imaging-derived tumor volumes, transcriptomic data, and serum biomarkers. The model could also be coupled with pharmacokinetic-pharmacodynamic (PK-PD) descriptions to study how targeted anti-EGFR therapies alter receptor activation over time. Together, these extensions may support the development of quantitative platforms for hypothesis generation, treatment stratification, and the design of more effective strategies to prevent or delay EGFR-driven tumor emergence.

## Data availability

The data and code used for calibration simulations of the 3D multiscale model are publicly available at https://github.com/Romasa/Cancer3D_validation/tree/main. The equations, parameter values, and numerical methods required to reproduce the continuous receptor-structured and reduced population-dynamics model results are provided in the manuscript.

## A Hybrid model calibration to physiological data

Across *in vivo* experiments, an EGF concentration from 0.45 - 1.63 ng/ml is indicated. In our simulation, we vary EGF concentrations from 0.05 to 1 ng/ml to include the cases of EGF scarcity. In the 3D multicellular model, discrete EGF molecules are introduced into the system. First, we calculate the number of EGF molecules by converting from ng/ml to the number of molecules using Avagadro’s number. The number of EGF molecules is then downsized by coarse-graining to reduce the computational cost. The molecular count of EGFs in the computational domain is calculated with a molecular weight for EGF of 6400 *g/mol* and *r* = 50 *µm*. Due to computational limitations, the model can consider up to 24 EGFR clusters per cell. This corresponds to approximately a one-hundredth of the real number of receptor clusters during EGFR overexpression [94]. We use the same ratio to reduce the number of EGF molecules. The number of EGF molecules that are considered for each physiological concentration is given in Table 5.

**Table 5.**
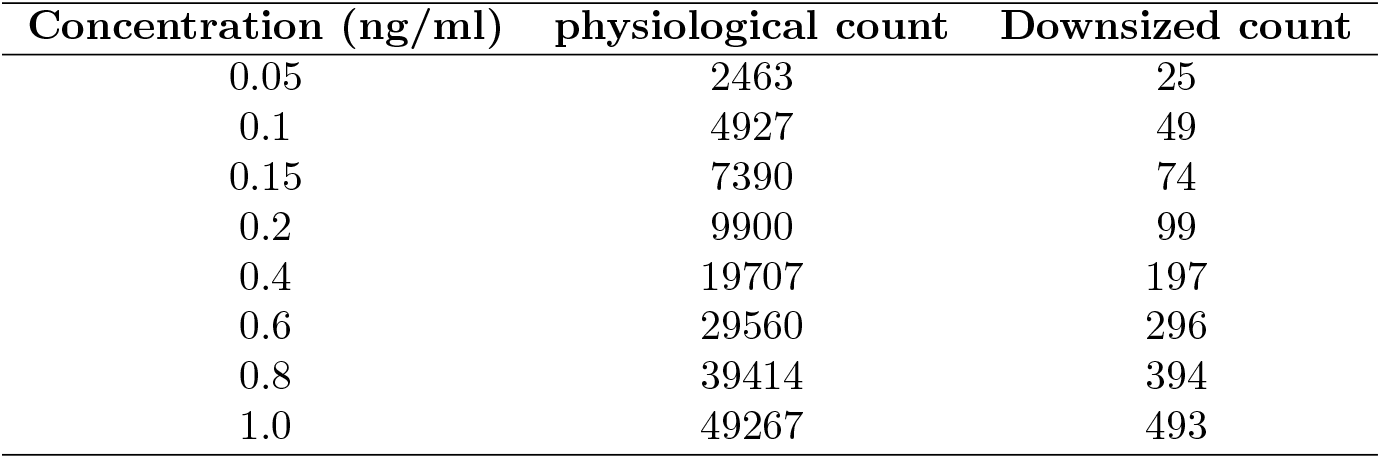
The conversion from EGF concentration to the number of molecules considered in the simulations.

## B Calibration of the intracellular ERK dynamics of the multiscale model

To reduce the computational time of simulations, the Brownian dynamics model for intracellular regulation in the original version of the model [61] was replaced by an ODE model. It describes the activation of ERK molecules, which depends on the number of active receptors, and their inactivation, modelled as an exponential decay:

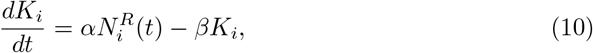

where *N*^*R*^(*t*) is the number of active receptors at any time *t, α* and *β* are positive constants. To determine their value, we use experiments by Orton et. al [74], where they incubated PC12 cells with EGF for a specified time interval to observe EGF stimulated ERK activation. The observed dynamics indicate active ERK level reached at 5-6 minutes, and started to decline from that point on through 40 minutes. Figure 10 shows that the simulated ERK concentration closely match the experimental data. The fitted values for the parameters *α* and *β* are given in Table 2.

**Fig 10.**
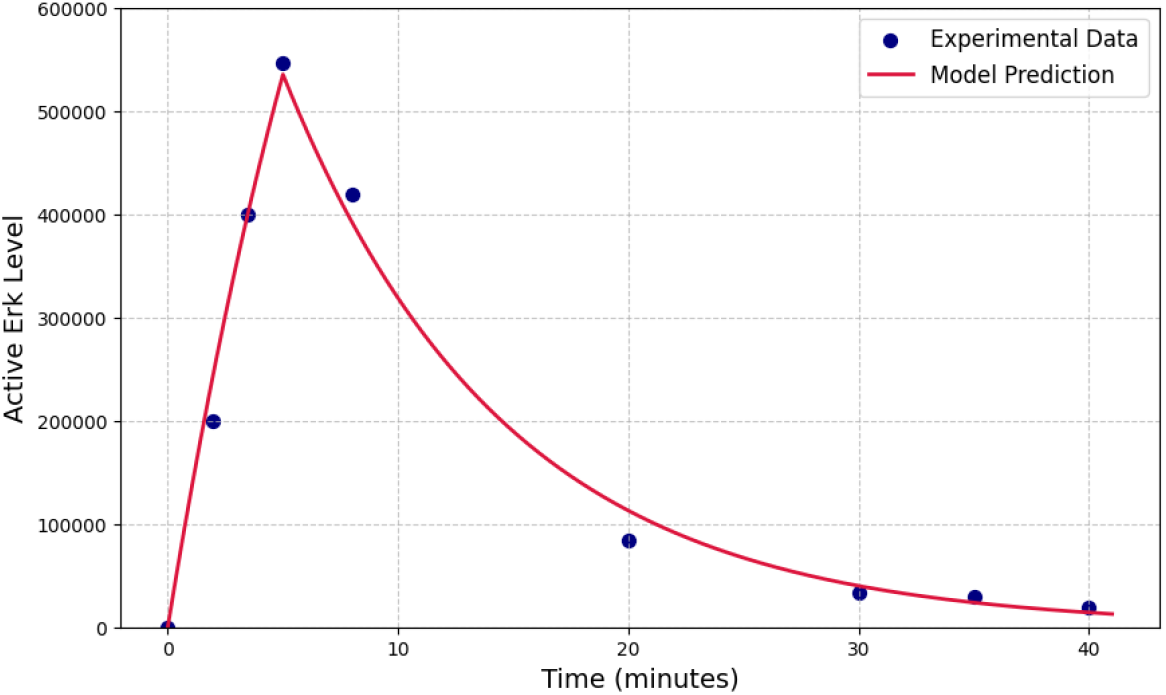
The results of simulations with the intracellular model for ERK levels following transient EGFR activation. Experimentally level shown as blue circles [74] were used to numerically fit the ODE model (Equation 10).

## C Stability analysis of the reduced system equilibrium points

This section presents a stability of the ODE system obtained from applying moment-reduction to the receptor-structured model. Let us consider the system:

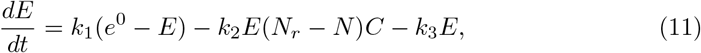

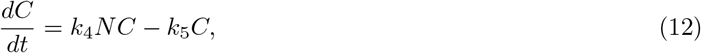

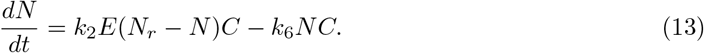

Let *F*_1_, *F*_2_, *F*_3_ denote the right-hand sides. The Jacobian matrix of the system is

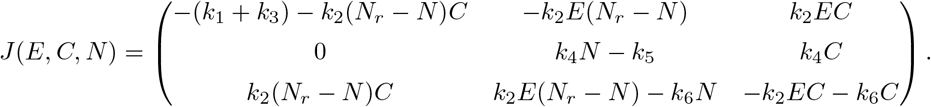

We analyze the stability of each equilibrium:

**Equilibrium 1:** 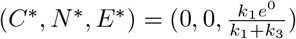 When evaluating the Jacobian at (0, 0, *E*\*), we obtain

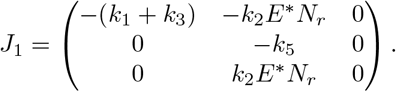

The eigenvalues satisfy

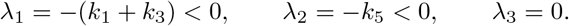

Thus Equilibrium 1 is non-hyperbolic, but stable in the *E*- and *C*-directions.

**Equilibrium 2:** *C*\* = 0, 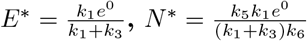

At this equilibrium we have *C*\* = 0 and *N* * > 0. The Jacobian becomes

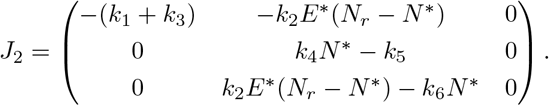

The characteristic polynomial reduces to

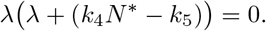

Thus

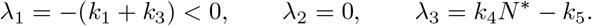

Hence Equilibrium 2 is stable if and only if

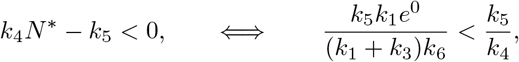

or equivalently

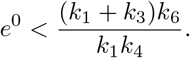

**Equilibrium 3:** 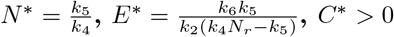

The third equilibrium satisfies

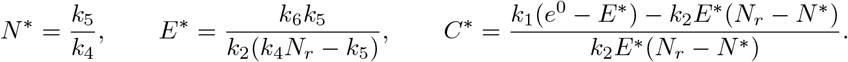

The equilibrium is biologically feasible only when *C*\* > 0, which requires

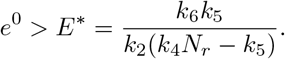

Linearisation yields one zero eigenvalue (due to *k*_4_*N* * *− k*_5_ = 0) and two remaining eigenvalues whose real parts are negative whenever *C*\* > 0. Thus Equilibrium 3 is stable whenever it exists, i.e. when

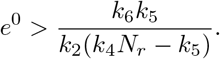

## D List of abbreviations

For convenience, we present a list of used abbreviations in Table 6.

**Table 6.** List of used abbreviations.

| Abbreviation | Meaning |
| --- | --- |
| EGF | epidermal growth factor |
| EGFR | epidermal growth factor receptor |
| CSC | cancer stem cell |
| RTK | receptor tyrosine kinase |
| TKI | tyrosine kinase inhibitor |
| MD | molecular dynamics |
| BD | Brownian dynamics |
| PK | pharmacokinetics |
| PD | pharmacodynamics |
| ODE | ordinary differential equation |
| PDE | partial differential equation |

